# The computational cost of explicit strategies in force-field adaptation

**DOI:** 10.64898/2026.11.24.768939

**Authors:** Hoi Kan, Yiyu Wang, Jordan A. Taylor

## Abstract

Sensorimotor adaptation is often attributed to an implicit, cerebellar-based process, particularly when learning to compensate for novel limb dynamics. However, recent evidence suggests that explicit strategies can also contribute to force-field adaptation, although how these strategies are represented and implemented is unclear. In visuomotor adaptation, explicit strategies can rely on an algorithmic-like process in which the time required to prepare a movement increases with the magnitude of the required correction. Here, we asked whether explicit compensation for force-field perturbations is generated through a similarly time-consuming computation. Human participants performed reaching movements in a paired-trial paradigm in which the force experienced on one movement could be used to prepare a counteracting response on the subsequent movement. Across two experiments, we found that when participants were encouraged to attempt to counteract the forces, movement preparation time increased systematically with force magnitude: larger compensatory responses required longer preparation. This relationship was substantially reduced when participants performed exactly the same task but were not informed about the paired-trial design, performance was significantly worse and was not accompanied by a corresponding increase in preparation time. These findings suggest that when participants use a strategy to improve performance in response to perturbing forces, it appears to be accomplished with a parametric, algorithmic-like process – one may be similar to strategies employed in visuomotor adaptation.

**Significance Statement:** Adapting to unexpected forces is often viewed as an implicit, cerebellar-dependent process, but recent work shows that explicit strategies can improve performance. How these strategies are represented and implemented remains unclear. Here, we asked whether force-field strategies rely on an algorithmic process, as observed in visuomotor adaptation. Across two experiments, explicit strategies improved compensation but came at a cost: movement preparation took longer as force magnitude increased. These findings suggest that strategic force compensation relies on a time-consuming algorithmic computation. Such computational demands may help explain why people avoid leveraging explicit strategies for force-field adaptation.

## Introduction

Humans routinely adjust their movements when interacting with unstable or unpredictable physical environments. This capacity for sensorimotor adaptation is evident when, for example, when a pianist encounters an unfamiliar instrument with a sticky key, they must quickly adapt by striking that key with greater force to produce the intended sound. More generally, sensorimotor adaptation allows the motor system to maintain successful performance when the mapping between motor commands and sensory feedback changes (Krakauer et al., 2019). This adaptation can reflect multiple processes, including implicit recalibration of motor commands as well as explicit strategies for counteracting a perturbation (Taylor et al., 2014, McDougle et al., 2015), but the relative contributions of these processes have been characterized quite differently across visuomotor and force-field adaptation paradigms. In visuomotor adaptation, learning reflects more of a balance between these processes, with explicit strategies rapidly improving performance through deliberate re-aiming of the movement plan, while implicit recalibration gradually updates the visuomotor mapping (Mazzoni & Krakauer, 2006; Taylor et al., 2014; McDougle et al., 2015). In contrast, forcefield adaptation has traditionally been characterized as predominantly implicit, with learning attributed to gradual, error-driven updates to an internal model of limb dynamics and relatively little contribution from explicit strategy (Shadmehr et al., 2010).

This view may stem, in part, from the strong implicit-learning phenotype of force-field adaptation, such as persistent aftereffects after the perturbation is removed (Shadmehr & Mussa-Ivaldi, 1994), trial-by-trial dependent changes in predictive force output (Thoroughman & Shadmehr, 2000), generalization patterns consistent with updating an internal model of limb dynamics (Gandolfo et al., 1996; Donchin et al., 2003; Thoroughman & Taylor, 2005), and that the cerebellum appears necessary for adaptation (Maschke et al., 2004; Smith & Shadmehr, 2005). Yet these features are not unique to force-field learning. Implicit recalibration during visuomotor adaptation shows many of the same behavioral signatures, even when isolated from explicit strategies (Morehead et al., 2017; Hutter & Taylor, 2018; Poh & Taylor, 2019). A more likely reason is that explicit strategies have simply been much harder to characterize in force-field tasks. In visuomotor adaptation, explicit strategy can often be expressed as a spatial aiming direction and measured directly through self-report (Taylor et al., 2014; Miyamoto et al., 2020). Force-field compensation, in contrast, is not naturally represented as a simple, spatial aiming location. Instead, successful compensation requires generating a force profile that depends on the experienced dynamics, making explicit contributions difficult to capture with standard report-based measures. This measurement challenge may have contributed to the longstanding view that explicit strategies play little role in force-field adaptation.

Despite these measurement challenges, recent work provides converging evidence that explicit strategies can contribute to force-field compensation, particularly when the task structure is relatively simple. Participants can report strategies during force-field learning, suggesting that explicit processes may contribute to force-field adaptation (McDougle et al., 2015). Moreover, participants can voluntarily suppress and reproduce aspects of their compensatory force output, indicating that at least part of force-field compensation is under explicit or voluntary control (Schween et al., 2020; Forano et al., 2021; Forano & Franklin, 2024). Finally, explicit instructions can impact force-field adaptation, although the learning outcomes are less stable (Zhou et al., 2022, Zhou et al., 2024).

These previous findings have only begun to establish a role for explicit strategies in force-field adaptation. Beyond demonstrating that some form of strategic control is possible, relatively little is known about how these strategies are represented or implemented. Work in visuomotor adaptation suggests that explicit re-aiming can rely on a time-consuming, algorithmic process when computing a solution. Critically, the time required to prepare the movement scales with the size of the visuomotor rotation: larger rotations incur greater preparation costs than smaller rotations, a hallmark of the parametric processing observed in classic mental rotation (Shepard & Metzler, 1971; Georgopoulos & Massey, 1987; Pellizzer & Georgopoulos, 1993; Bhat & Sanes, 1998; McDougle & Taylor, 2019).

Building on these findings, we ask whether explicit strategies for force-field adaptation are generated through a similarly algorithmic process. Here, we used a paired-trial design, similar to that used to establish algorithmic strategy use in visuomotor adaptation (Velázquez-Vargas & Taylor, 2024, McDougle & Taylor, 2019), to test whether participants could use the force experienced on one movement to prepare a counteracting response on the next and if this comes at a preparation time cost. If participants leverage an algorithmic strategy to compute a force-specific counteracting response, then preparation time should scale with force magnitude.

## Materials and Methods

### Participants

Participants were recruited from the study subject pool maintained by the Department of Psychology at Princeton University and were provided with course credit or $20/hour in exchange for their participation. The study protocol was approved by the Institutional Review Board at Princeton University, and all participants provided written, informed consent prior to participation.

Across two experiments, 77 participants were randomly assigned into either experiment group or control group in each experiment. In experiment 1, 20 participants were assigned to the Explicit group (9 reported female, 9 reported male; mean age = 22.7 ± 5.6 years, range = 18– 40) and 20 participants to the Control group (15 female, 5 male; mean age = 20.8 ± 3.5 years, range = 18–30). All participants reported being right-handed according to the Edinburgh Handedness Inventory (Oldfield 1971). In experiment 2, 21 participants completed the Explicit condition (6 female, 15 male; mean age = 19.7 ± 1.19 years, range = 18–22) and 16 completed the Control condition (9 female, 7 male; mean age = 20.9 ± 5.48 years, range = 18–41). One participant in the Explicit condition and two participants in the Control condition reported being left-handed, but they still used their right hand for the experiment.

### Apparatus

Participants performed center-out reaching movements using a KINARM Endpoint robotic manipulandum (KINARM, Kingston, Ontario, Canada) controlled using Dexterit-E 3.8.2. Participants were seated in front of the robot and held the manipulandum handle with their right hand. All visual stimuli were presented onto a horizontally mounted mirror by a LCD monitor mounted vertically 22 cm above the mirror. Because the robotic manipulandum was 22 cm below the mirror, it creates the illusion that the cursor and targets are in the same horizontal plane of the movements. The cursor was 0.2 cm in diameter, the central start position was 0.7 cm in diameter, and each target was 0.7 cm in diameter. Kinematic and force data were sampled at 1,000 Hz.

### Experiment 1 Task Procedure

For participants assigned to the Explicit condition, a trial began with the manipulandum gently guiding the hand to the central starting position. After holding this position for 1250 ± 750 ms in the first trial (Observation) or 450 ± 150 ms in the second trial (Counteract), a target would appear 10 cm away from the start position either at 90° or -90° (Fig. 1B). Participants were instructed to make a rapid, ballistic shooting movement through the target rather than stopping within it. They could remain at the start position for as long as needed before initiating the movement, but were instructed to complete the reach rapidly once the movement began. Participants were required to complete the reaching movement within 250 ms, which was measured as the time it took for the hand to pass through the 10 cm radial distance to the target after leaving the central start position’s boundary. Once the cursor passed the 10cm radial distance to the target, the center, target and cursor disappeared. If the movement time exceeded this 250 ms limit, then the message “Too Slow” was presented on the visual display. This time constraint was intended to discourage slow movements and the contribution of online feedback corrections. After a 200ms delay following movement completion, the manipulandum automatically returned the hand to the central start position to begin the next trial.

**Figure 1.**
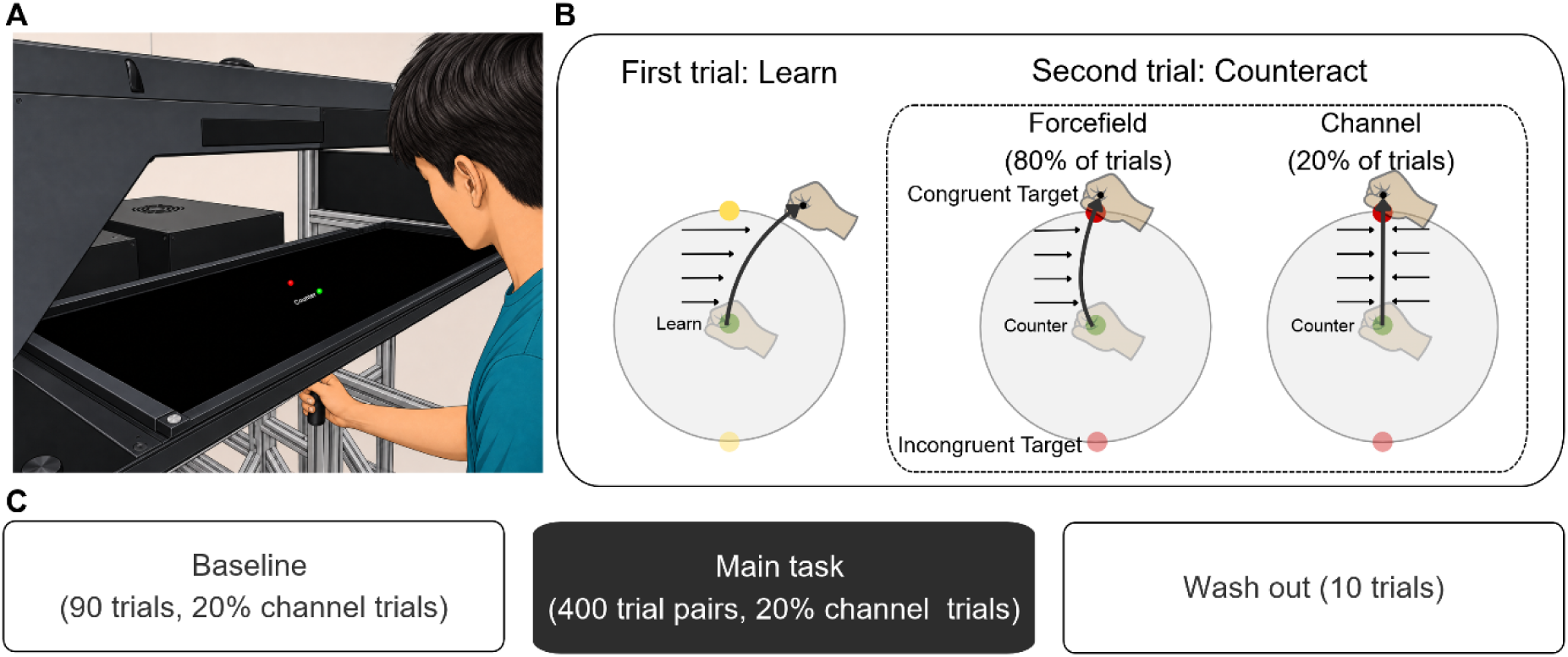
Experiment setup. ***A***, Participants used their right hand to control a visually displayed cursor via a robotic manipulandum. The visual workspace was projected onto a horizontally mounted mirror that blocked direct vision of the hand. ***B***, Trial pair schedule. The main task consisted of paired Observation and Counteract trials. On each pair, the Counteract trial target either repeated the Learn trial target location, producing a congruent target pair, or appeared at the opposite target location, producing an incongruent target pair. The figure shows an example of a clockwise force-field. ***C***, Experiment timeline. Participants completed a Baseline block, the main paired trial task, and a final force-channel probe block.

The primary task unfolded in a series of trial pairs (Fig. 1B). The first trial of each pair was a Learn trial, during which participants were instructed to reach directly to the target while experiencing a velocity dependent curl force-field. The magnitude of the applied force was proportional to hand velocity, and its direction was perpendicular to the direction of movement as follows:

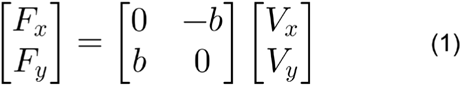

Where *F_x_* and *F_y_* are forces applied at the hand, *V_x_* and *V_y_* are hand velocities, and *b* determines the magnitude and direction of the force-field. The force-field magnitude was either 0, +/-7.5, or +/-15 N·s/m, and force direction was clockwise or counterclockwise (Eqn. 1), which was pseudorandomized and counterbalanced so that each magnitude and direction combination occurred equally often. Learn trials were indicated by the message “Learn” displayed on the screen and the target was in yellow. If the movement exceeded the 250 ms time limit, “Too slow” was displayed instead. After movement completion, the manipulandum gently returned the hand to the central start position to begin the next trial of the pair.

The second trial of each pair was a “Counter” trial, in which participants were instructed to counteract the force experienced on the preceding Learn trial. On 50% of the trials, the target could appear at the same target location as on the Learn trial (Congruent trials) or, on the other half, it could appear 180° away (Incongruent trials). This was to prevent participants planning the movement before they see the target, half the time the target appears at the same location during the Learn and Counteract trials (Congruent Trials) and half the time the target appears at the opposite locations from Learn to Counteract trials (Incongruent Trials) (Figure 1B). 80% of the Counteract trials are non-channel trials (Counteract trials). In those trials, the force-field was identical to that experienced on the preceding Learn trial. On 20% of the Counteract trials, a force channel was applied (Counteract-channel trials). The force channel trials were designed to obtain an accurate measure of the force exerted by the participants by restricting lateral hand deviations with a simulated spring with a stiffness of 4000 N/m and a damper of 10 N·s/m oriented perpendicular to the target direction (McDougle, Bond & Taylor, 2015; Scheidt et al. 2000; Smith et al. 2006). The occurrence of these channel trials was pseudorandomized across trials such that all force magnitudes, directions, congruent and incongruent trials were experienced equally as often. To encourage participants to counteract the force-field on Counteract trials, they were provided with coarse, binary performance feedback: “Good” if the cursor hit the target and “Miss” otherwise. Note, no performance feedback was provided on Learn trials.

To help participants notice the trial pair sequence, in between each trial pair, a 500-ms delay was imposed and the message “Relax” was visually displayed. Target onset was jittered to reduce anticipatory or rhythmic responding. On Learn trials, the target appeared 1,250 ± 750 ms after the cursor was stably positioned at the start location. On Counteract trials, the target appeared after 450 ± 150 ms. On Learn trials, the only possible feedback was “Too Slow”.

Participants first completed a brief tutorial to familiarize themselves with the reaching task. They then completed 90 trials without force-field perturbations (Baseline block). The targets could appear at 90° or -90° with equal likelihood. Force channels were randomly imposed on 20% of these trials to obtain a measure of baseline biases prior to introduction of the force-fields. Following the Baseline block, participants completed a primary Test block consisting of 800 trials arranged into 400 Learn and Counteract trial pairs. At the end of the Test block, participants completed 10 additional force-channel trials to measure whether any residual compensatory forces remained at the end of the session (Washout block).

The Control condition followed the same trial sequencing and block structure as for the Explicit condition, but importantly, they were not informed of the paired-trial sequence and the messages of “Learn” and “Counter” were not displayed. In addition, the between trial pair “Relax” message was also removed. All intertrial intervals were drawn from the same intertrial distribution (450 ± 150-ms) to make the pairing between successive trials less apparent. As such, there was virtually no cue as to the identity of the trials.

### Experiment 2 Task Procedure

Experiment 2 tested whether the relationship between preparation time and force compensation persisted for point-to-point reaches, in which participants were required to terminate their movement at the target, as is more conventional in force-field adaptation studies. A trial ended once the center of the cursor stopped within the target radius and hand velocity fell below 2.5 cm/s. Participants were required to complete the movement within 400–700 ms. Movements completed in less than 400 ms elicited the message “Too Fast,” whereas movements exceeding 700 ms elicited the message “Too Slow.” In addition, because Counteract trials continued until the cursor terminated at the target, “Miss” feedback was not used. Aside from these changes to how a trial unfolded, the paired-trial sequencing and block-level structure were identical to Experiment 1. The Explicit and Control groups received otherwise identical task instructions, except that participants in the Explicit group were informed about the paired-trial sequence, whereas participants in the Control group received no information about this structure.

### Movement analysis

Our primary analysis asked whether the preparation time scaled with the force generated on all Counteract trials, which would be indicative of an algorithmic-like strategy that has been observed in studies of visuomotor adaptation (McDougle & Taylor 2019). We first assessed the force generation using two complementary measures of counteracting force production. For our first measure, we examined the Counteract trials performed in the force-field (80% of trials) where lateral hand position provides a kinematic measure of how effectively participants counteracted the perturbation experienced on the preceding Learn trial. For both experiments, lateral displacement was measured at peak velocity. We then assessed the relationship between force perturbation magnitude in the Learn trial and the preparation time cost in the Counteract trial.

Because lateral position can reflect the combined influence of predictive force generation, the imposed force-field, and online corrective movements, for our second measure, we examined the remaining 20% of Counteract trials, in which movements were performed in a force channel. Because the channel constrained the hand to a straight path toward the target, lateral force exerted against the channel could be measured with minimal lateral displacement or contribution from online corrective responses. For each force-channel trial, lateral force was defined relative to the straight-line movement axis. Positive values indicated force exerted toward the left. Force channel recordings were baseline corrected for each participant based on their channel force profile during the null force Baseline trials, so that any bias in lateral force for upward and downward movements toward 90° and 270° would cancel out in analyses of Counteract-channel trials. The force channel recordings were compared to the perturbation force magnitude in the previous Learn trial, with the signs adjusted to represent the correct counteracting direction in the current channel trial.

For both experiments, the lateral force was defined as the analyzed from the lateral force at peak velocity of the movement. For each trial, the measured lateral-force profile was compared with the ideal force profile required to counteract the force-field experienced on the preceding Learn trial. The ideal force profile was calculated from the participant’s hand velocity and the imposed force-field magnitude and direction from the preceding Learn trial. An adaptation index was obtained by regressing the measured lateral force profile onto the ideal force profile, which is defined as the lateral force the participant would need to generate over time to perfectly counteract the velocity dependent force-field, with the intercept constrained to zero (Coltman et al., 2019). An index of 1 indicated complete compensation for the preceding force-field, an index over 1 indicates overcompensation, an index of 0 indicated no systematic compensation, and a negative value indicated force production in the incorrect direction.

To further minimize online corrective responses, Counteract trials and Counteract-channel trials were restricted to movement times that fell within the task constraints. In Experiment 1, in which participants shot through the target, we included only trials with movement durations under 250 ms (98.92% in the Explicit group; 97.24% in the Control group). In Experiment 2, in which participants stopped at the target, force output trials were retained if movement time (MT) fell within ±2.5 standard deviations of the mean (97.37% in the Explicit group; 97.66% in the Control group).

Preparation time was defined as the interval between target onset and movement onset, with movement onset identified as the moment the cursor first left the start-position boundary (0.7 cm in diameter). Movement time was defined as the interval between movement onset and the cursor reaching a radial distance of 10 cm in Experiment 1, and between movement onset and the cursor entering the target with hand speed below 2.5 cm/s in Experiment 2. Practice and tutorial trials, and Washout trials were excluded from the main analyses.

### Statistical analysis

We hypothesized that the (1) adaptation index should remain unchanged over the course of the experiment. We further hypothesized that (2) counteracting force would scale with the direction and magnitude of the observed force in the Explicit group, and that (3) this scaling would be stronger in the Explicit group than in the Control group. Similarly, we hypothesized that (4) preparation time would increase with the magnitude of the force participants need to counteract for the Explicit group, and that (5) this preparation time scaling would also be stronger in the Explicit group than in the Control group.

To test the first hypothesis, two simple linear regressions were performed for the Explicit and Control group. The adaptation index was regressed against the trial count across the course of the experiment.

To test the second and third hypotheses, a simple linear regression was fit for each participant, with lateral force magnitude during Counteract-channel trials as the outcome variable and the force experienced during the preceding Learn trial (-15, 7.5, 0, 7.5, or 15 N·s/m) as the predictor, with positive values representing rightward force (Forcefield in CW upward movement and CCW downward movement.), and negative values representing leftward force (Forcefield in CW downward movement and CCW upward movement.). The resulting regression slopes were compared with zero separately for the Explicit and Control groups using one-sample t-tests and were then compared between groups using an independent-samples t-test. Since channel trials only account for 20% of the Counteract trials, the remaining 80% counteract forcefield trials were analyzed in terms of lateral displacement change. If the correct counteracting force is applied to the forcefield, the lateral displacement at peak velocity would be lower than the displacement in the Learn trial. A simple linear regression was fit for each participant, with lateral displacement at peak velocity during counteracting forcefield trials as the outcome variable and the force experienced as the predictor. The regression slopes were compared with zero for Explicit and Control groups using one-sample t-tests and compared between groups using independent-sample t-test.

To test the fourth and fifth hypotheses, a simple linear regression was fit for each participant, with preparation time as the outcome variable and the force perturbation magnitude in the Learn trial as the predictor. The resulting regression slope quantified the change in preparation time associated with increasing force perturbation. Within each group, slopes were compared with zero using separate one-sample t-tests. A positive slope significantly greater than zero would indicate that preparation time increased parametrically with force magnitude. Slopes were then compared between the Explicit and Control groups using an independent samples t-test to determine whether force dependent preparation time scaling differed between groups.

In an exploratory analysis, we compared the difference in the preparation time scaling effect between congruent and incongruent targets on preparation time in the Explicit group by fitting separate linear regressions to congruent and incongruent channel Counteract trials. Slopes for both kinds of trial pairs were compared with zero using one-sample t-tests, and then compared with each other to determine if they are truly different using paired sample t-tests.

## Results

### Experiment 1

#### Counteracting force scaled with observed force within trial pairs

The goal of the present work was to determine whether the preparation time required to use an explicit strategy to counteract a force perturbation scales with perturbation magnitude. Such scaling would suggest that explicit strategies for force production rely on an algorithmic-like process similar to that observed in visuomotor rotation tasks. Participants were exposed to force perturbations of varying magnitudes and directions in a trial pair design where the first trial of the pair (Learn trial) they were instructed to reach directly to the target to experience the perturbation; in the second trial of the pair (Counteract trial), participants in the Explicit group were instructed to counteract the force they just experienced, while participants in the Control group were not informed of the trial pair design.

Prior to the introduction of the forces, participants performed a short Baseline block to familiarize themselves with the task. At the end of this block, there were no significant differences between the Explicit and Control groups in terms of lateral displacement at peak velocity (t(38) = -0.89, *p* = 0.38), lateral force production at peak velocity (t(38) = -0.95, *p* = 0.35), movement time or preparation time (*p*’s > 0.05). Following the Baseline block, the Explicit group received additional instructions regarding the trial pair design.

As expected, lateral displacement scaled with the applied force perturbation on the Learn trials for both the Explicit and Control groups (*p*’s < 0.05; Figure 2A). To ensure participants, at least in the Explicit group, were attempting to counteract the perturbation they experienced during the Learn trial, we examined their behavior on the subsequent trial. These subsequent trials could take on two forms, either without a force channel (Counteract trial) or with a force channel (Counteract-channel trial). If lateral displacement is lower in the Counteract trial than the Learn trial, it would indicate that the Explicit group followed the instructions to attempt to counteract the force. Similarly, if the lateral force in the Counteract-channel trials scaled with the force applied on the Learn trial, this would also indicate that they were attempting to appropriately counteract the force.

**Figure 2.**
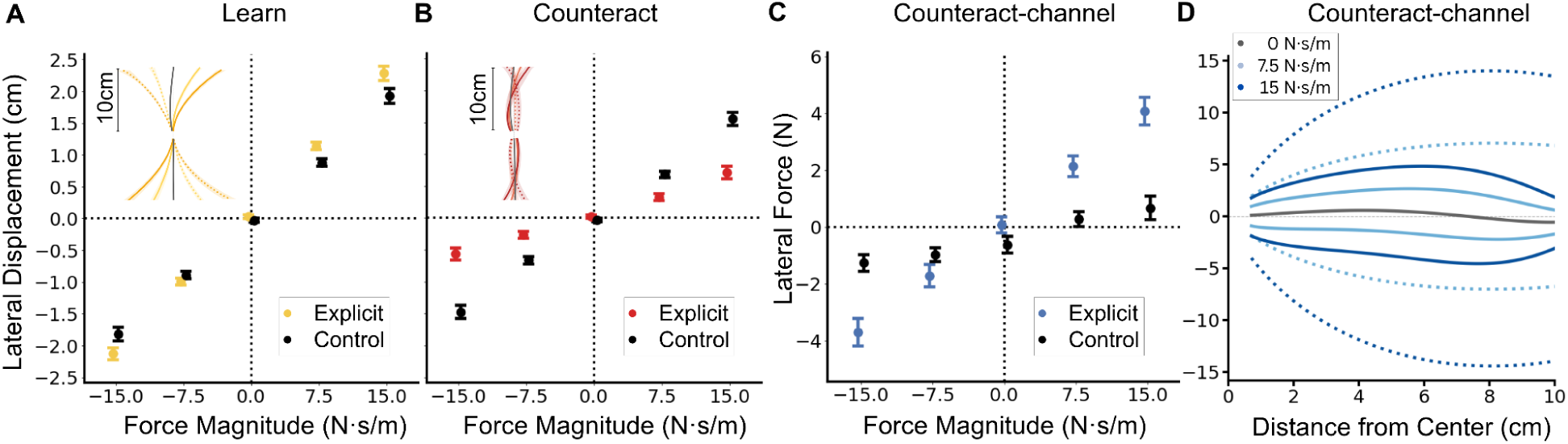
Experiment 1 task performance. ***A***, Learn trial lateral displacement. The main panel shows the lateral displacement at peak velocity as a function of perturbation magnitude. The inset shows mean Learn trial trajectories: dark, 15 N·s/m; light, 7.5 N·s/m; gray, no force. Solid and dotted lines indicate CW and CCW force-fields, respectively. ***B***, Counteract trial lateral displacement (no force channel). The main panel shows the lateral displacement at peak velocity as a function of perturbation magnitude. The inset shows mean Counteract trial trajectories. ***C***, Counteract-channel trial lateral counteracting force at peak velocity as a function of force magnitudes (force channel applied). ***D***, Channel Counteract trial force profiles. Dotted lines stand for ideal force profile and solid lines stand for recorded force profile.

Indeed, we observed a clear reduction in the lateral displacement on Counteract trials (Figure 2A, B). Both the Explicit and Control groups exhibited reduced deviation during Counteract trials compared with Learn trials (*p*’s < 0.001), as would be expected based on previous reports of single-trial based force adaptation (Fine & Thoroughman, 2006). However, we observed a greater reduction in lateral displacement for the Explicit group compared to the Control group (*p* < 0.001; Figure 2A,B), suggesting that they may have leveraged an explicit strategy to counteract the forces as instructed.

Counteract-channel trials provided a complementary way to assess whether participants in the Explicit group used a strategy to counteract the preceding perturbation (Figure 2C). For the Explicit group, lateral force at peak velocity scaled proportionally with the observed force, increasing by 0.25 N for every 1 N·s/m increase in perturbation strength (t(19) = 8.87, *p* < 0.001).

For the Control group, lateral force at peak velocity also scaled with the observed force, but to a lesser extent, increasing by 0.10 N for every 1 N·s/m increase (t(19) = 5.35, *p* < 0.001). Indeed, the slopes were lower for the Control group than for the Explicit group (t(38) = 4.62, *p* < 0.001), suggesting that the Explicit group may have deliberately attempted to counteract the forces experienced in the Learn trial – a hallmark of applying an explicit strategy. These lateral deviation and lateral force findings provide converging evidence that participants are more accurate when they were informed of the task structure and were instructed to use a strategy to counteract the forces.

To prevent accumulation of adaptation over the course of the experiment, we randomized the trial-by-trial force-field (Scheidt et al. 2001; Fine & Thoroughman 2006). We nevertheless examined whether counteracting force production improved over the course of the experiment. The adaptation index did not change across trials for both the Control group (t(19) = -0.27, *p* = 0.79), nor the Explicit group (t(19) = −0.62, *p* = 0.55), providing no evidence of a gradual buildup of adaptation or improvement in applying an explicit strategy to counteract the forces. An additional confirmation was gleaned from the Washout block where a force channel was imposed at both targets and participants were instructed to reach directly to the target. There was no significant change in force production from Baseline force channel trials for either the Explicit group (t(19) = -0.83, *p* = 0.42) and for the Control group (t(19) = -1.55, *p* = 0.14). No significant difference between groups (t(38) = -1.35, *p* = 0.19). Movement time did not differ significantly between groups (*p* = .84) while preparation times were 805 ms longer for the Explicit group than the Control group (*p* < .001), possibly because participants in the Explicit group had grown accustomed to the trial pair structure used earlier in the experiment.

### Preparation time-force profile indicates the recruitment of algorithmic strategy

While the difference in lateral displacement and force production between the Explicit and Control group suggests use of a strategy, a second indication would be whether the Explicit group displayed a preparation time increase on Counteract trials. If the preparation time also scales with the magnitude of the force to be generated, it would further suggest that strategies for force adaptation could be the outcome of an algorithmic-like process. To test this, we combined the preparation times for Counteract and Counteract-channel trials since participants could not know the trial type in advance. For the Explicit group, we found a significant increase in preparation time on Counteract trials relative to the Learn trials (t(19) = 4.39, *p* < 0.001; Figure 3A,B). Furthermore, for the Explicit group, a simple linear regression revealed that preparation time scaled with force magnitude, having a mean of 41.6 ms for every 1 N·s/m increase (t(19) = 8.47, *p* < 0.001). In contrast, for the Control group, preparation time on Counteract trials did not increase relative to the Learn trial (t(19) = 1.46, p = 0.16; Figure 3A,B) and, correspondingly, the slope for the Control group did not differ from zero (t(19) = -0.08, p = 0.93). A paired sample t-test of the regression slopes revealed a significant difference between groups (t(38) = 8.26, p < 0.001) with preparation time scaling more strongly with force production in the Explicit group than in the Control group (Figure 3A). There was no relationship between force magnitude and preparation on Learn trials for both the Explicit and Control groups (*p*’s > 0.05; Figure 3A).

**Figure 3.**
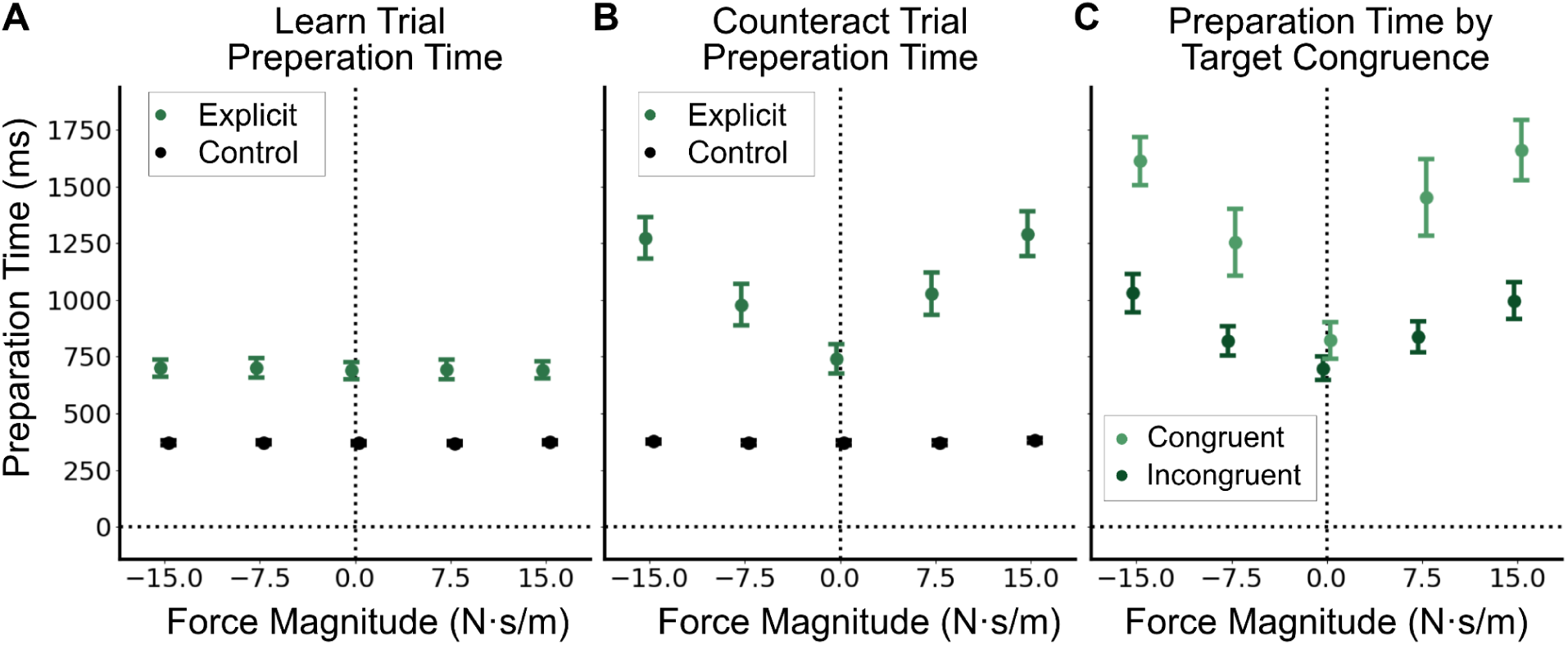
Experiment 1 preparation time cost. ***A***, Learn trial Preparation time vs. force magnitudes between the Explicit and Control group. ***B***, Counteract trial Preparation time vs. force magnitudes between the Explicit and Control group. ***C***, Preparation time vs. force magnitudes for the Explicit group, between Congruent and Incongruent target locations.

To prevent participants from preparing the movement before the target onset, the target could appear in the same or different location within a trial pair. We anticipated that when the target appeared at a different location (incongruent trials), there would be a constant increase in preparation time across the force magnitudes, as a sign flip would be needed for the counteracting direction. Interestingly, we found not only an increase in the intercept (t(19) = - 3.20, *p* < 0.05) but also an increase in the slope for incongruent trials compared to congruent trials (t(19) = 5.89, *p* < 0.001; Figure 3B). While congruent trials have a slope of 25.99 ms/N (t(19) = 5.67, *p* < 0.001), incongruent trials have a slope of 57.41ms/N (t(19) = 8.92, p < 0.001).

### Experiment 2

#### Counteracting force scaled with observed force within trial pairs

Experiment 2 was designed to test whether parametric preparation time scaling generalizes to the Stop-at-Target paradigm, which is more commonly used in force-field experiments. We find that this paradigm paints a similar picture to the Shoot-Through-Target paradigm (Experiment 1).

At the end of the Baseline block, prior to introduction of the force perturbation, there were no significant differences between the Explicit and Control groups in terms of lateral displacement at peak velocity (t(35) = 1.34, *p* = 0.19), force production on channel trials (t(35) = -0.11, *p* = 0.29), nor movement time or preparation time (*p*’s > 0.05).

As expected, lateral displacement scaled with the applied force perturbation on the Learn trials (*p* < 0.05; Figure 4A). As in Experiment 1, we observed a clear reduction in the lateral displacement on Counteract trials compared with Learn trials for both the Explicit and Control groups (*p*’s < 0.001; Figure 4A, B). However, we observed a greater reduction in lateral displacement for the Explicit group compared to the Control group (*p* < 0.001; Figure 4A, B)

**Figure 4.**
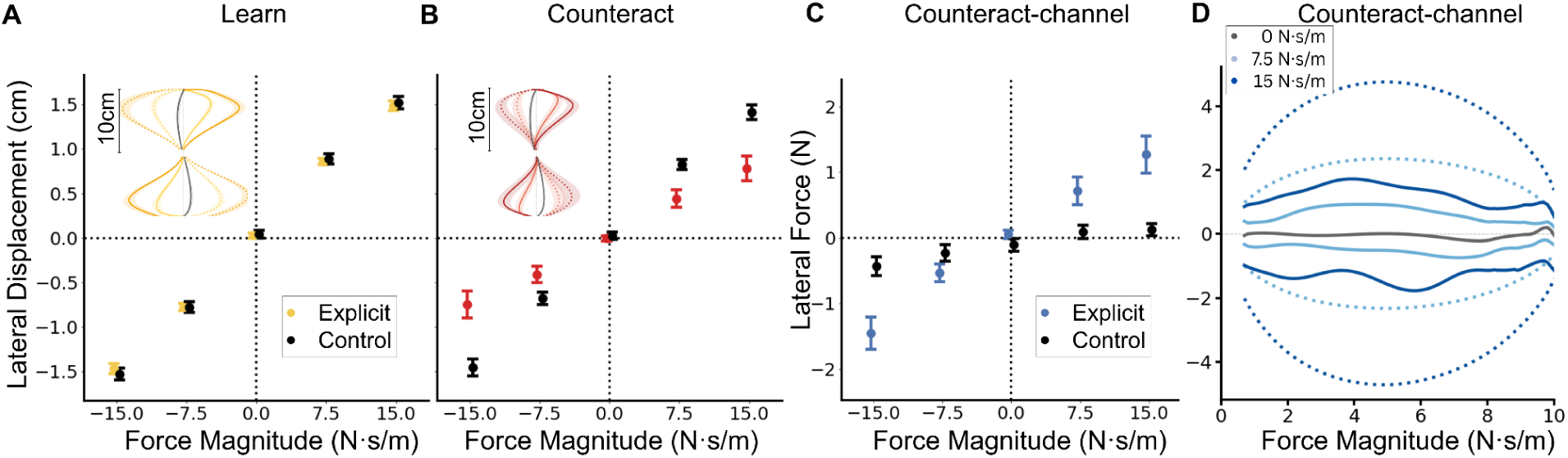
Experiment 2 task performance. ***A***, Learn trial lateral displacement. The main panel shows the lateral displacement at peak velocity as a function of perturbation magnitude. The inset shows mean Learn trial trajectories: dark, 15 N·s/m; light, 7.5 N·s/m; gray, no force. Solid and dotted lines indicate CW and CCW force-fields, respectively. ***B***, Counteract trial lateral displacement (no force channel). The main panel shows the lateral displacement at peak velocity as a function of perturbation magnitude. The inset shows mean Counteract trial trajectories. ***C***, Counteract-channel trial lateral counteracting force at peak velocity as a function of force magnitudes (force channel applied). ***D***, Channel Counteract trial force profiles. Dotted lines stand for ideal force profile and solid lines stand for recorded force profile.

For the Counteract-channel trials from the Explicit group, lateral force at peak velocity scaled proportionally with the observed force, increasing by 0.09 N for every 1 N·s/m increase in perturbation strength (t(20) = 5.49, *p* < 0.001). For the Control group, lateral force at peak velocity also scaled with the observed force, but to a lesser extent, increasing by 0.02 N for every 1 N·s/m increase (t(15) = 4.65, *p* < 0.001). However, the slopes were lower for the Control group than for the Explicit group (t(35) = 4.18, *p* < 0.001). Taken together, the greater reduction in lateral displacement and greater force production for the Explicit group suggests that they may have been using a strategy to counteract the force perturbations, consistent with the findings of Experiment 1.

To confirm that adaptation did not accumulate over the course of training, we analyzed both the adaptation index during the Test block and force production on channel trials in the Washout block. As in Experiment 1, the adaptation index did not change over the course of the experiment for the Explicit group (t(20) = -2.01, *p* = 0.06); however, the adaptation index increased modestly by ∼0.002 every trial (t(15) = 3.49, *p* < 0.05) for the Control group. During the Washout block, there was also a small yet significant change in force production compared to the Baseline block for both the Explicit group (0.1N; t(20) = -3.02, *p* < 0.05) and the Control group (0.2N; t(16) = 2.87, *p* < 0.05). Importantly, however, this was not significant between groups (t(35) = -0.21, *p* = 0.83). Movement time for the Control group is 47ms higher than the Explicit group (*p* < 0.05) while preparation times were 477ms longer for the Explicit group than the Control group (*p* < .001), possibly because participants in the Explicit group had grown accustomed to the trial pair structure used earlier in the experiment.

#### Preparation time-force profile indicates the recruitment of algorithmic strategy

Preparation times for Counteract and Counteract-channel trials were examined for signatures of strategy use. For the Explicit group, we found a significant increase in preparation time on Counteract trials relative to the Learn trials (t(20) = 4.74, *p* < 0.001; Figure 5A,B). Furthermore, for the Explicit group, a simple linear regression revealed that preparation time scaled with force magnitude, having a mean of 31.49 ms for every 1 N·s/m increase (t(20) = 6.28, *p* < 0.001). In contrast, for the Control group, preparation time on Counteract trials were not different from Learn trials (t(15) = 1.15, p = 0.27; Figure 5A,B)and, correspondingly, the slope for the Control group did not differ from zero (t(15) = 1.59, *p* = 0.13). A paired sample t-test of the regression slopes revealed a significant difference between groups (t(35) = 5.21, *p* < 0.001) with preparation time scaling more strongly with force production in the Explicit group than in the Control group (Figure 5A). There was no relationship between force magnitude and preparation on Learn trials for both the Explicit and Control groups (*p*’s > 0.05).

**Figure 5.**
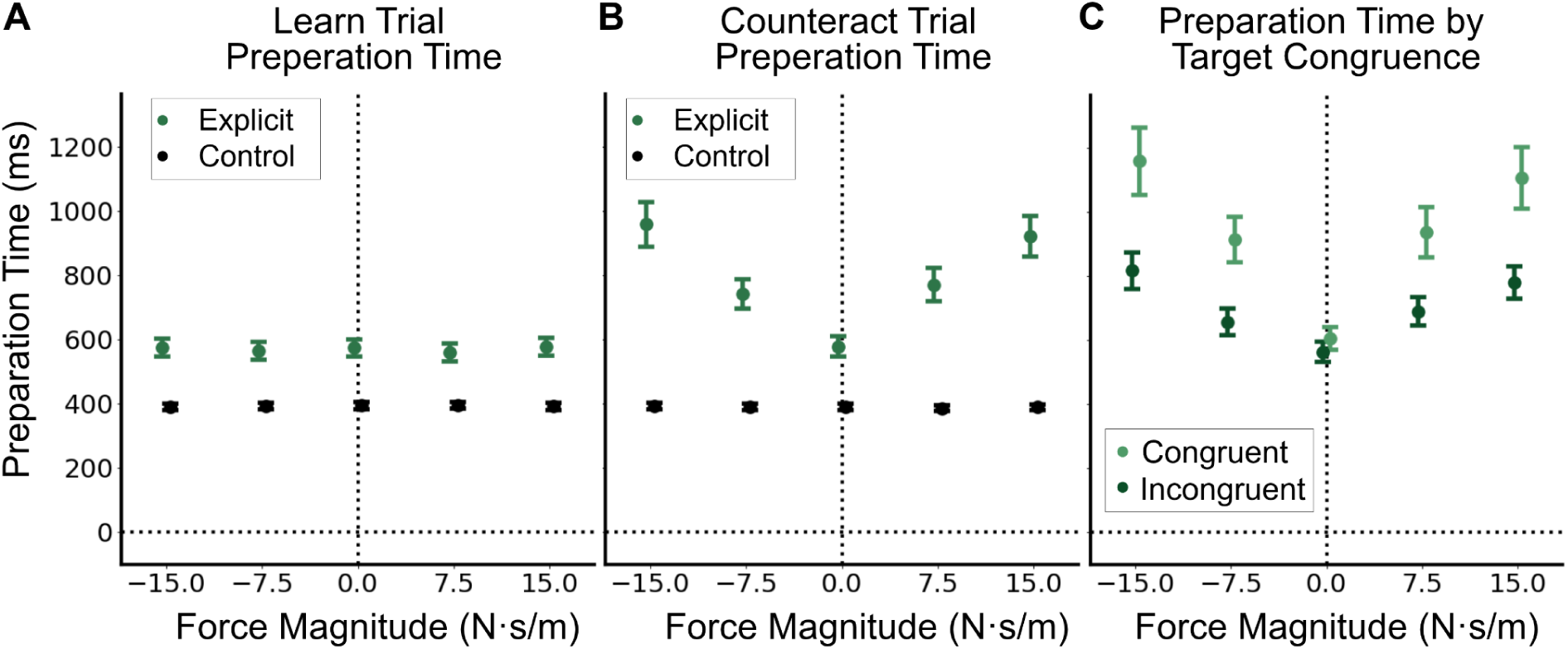
Experiment 2 preparation time cost. ***A***, Learn trial Preparation time vs. force magnitudes between the Explicit and Control group. ***B***, Counteract trial Preparation time vs. force magnitudes between the Explicit and Control group. ***C***, Preparation time vs. force magnitudes in the Explicit group, between Congruent and Incongruent target locations.

To prevent participants from pre-planning their movements, the target could change location between Learn and Counteract trials. As in Experiment 1, we found a target congruence effect: preparation time was slower (t(20) = -3.26, p < 0.05) and steeper (t(20) = -4.11, p < 0.001) for incongruent targets for the Explicit group ((Figure 5B). Congruent trials had a slope of 22.70 ms/N (t(20) = 5.63, p < 0.001), while incongruent trials had a slope of 40.40 ms/N (t(20) = 6.14, p < 0.001).

## Discussion

To gain better insight into how force adaptation strategies may be implemented and represented, we tested whether they rely on an algorithmic-like process similar to that observed in visuomotor adaptation (Georgopoulos & Massey, 1987; Pellizzer & Georgopoulos, 1993; Bhat & Sanes, 1998; McDougle & Taylor, 2019). To test this, we used a trial pair structure in which participants first experienced a force perturbation of varying magnitude and then attempted to counteract it on the subsequent trial. We found that preparation time proportionally increased with force magnitude, suggesting that force adaptation can recruit an algorithmic-like strategy.

Converging evidence from a wide range of studies has supported the traditional view that force-field adaptation reflects the gradual updating of an internal model through a cerebellar-dependent, implicit and automatic process (Shadmehr & Krakauer, 2010; Krakauer et al., 2019). However, more recent work suggests that explicit strategies can speed learning under certain situations, particularly when instructions are provided (Zhou et al. 2022, 2024) or the adaptation problem is relatively simple (McDougle et al. 2015; Schween et al., 2020; Forano et al., 2021). Their contribution to adaptation may be greatest early in training and diminish as implicit adaptation accumulates. To encourage persistent strategy use, we limited the accumulation of implicit adaptation by randomizing the perturbations between the “observe-then-counteract” trial pairs and informed the Explicit group of the trial pair structure (Scheidt et al., 2001; Fine & Thoroughman, 2006). We found that the Explicit group displayed a greater reduction in error on Counteract trials compared to the Control group who was not informed of the trial structure. This was not a small difference but a 26% reduction in error from just a single trial.

However, this substantial performance benefit of using a strategy to counteract a force-field came with a corresponding cost in preparation time. The observed error reductions of 49% in Experiment 1 and 20% in Experiment 2 were accompanied by increases in preparation time of 128% and 71%, respectively, with even larger increases when the target switched location. Although these preparation time costs amounted to only fractions of a second on each trial, even small increases in cognitive demand may influence whether people choose to use a strategy (Kool et al., 2010; Shenhav et al., 2013). Consistent with this possibility, we recently found that participants were sensitive to subtle differences in the demands of implementing a visuomotor mental rotation strategy and tended to avoid the more demanding option when given a choice (Kim et al., 2024). The preparation time cost observed here was also considerably greater than that reported for visuomotor rotation. Here, we found a preparation time increase of approximately 24 ms for every degree of lateral displacement reduction, compared with only 4.4 ms per degree in a visuomotor rotation task (McDougle & Taylor, 2019). While these measures may not perfectly capture the computational demands between the tasks, the difference suggests that strategic compensation for force-fields may be much more costly. Such costs could discourage strategy use when the incentive to restore performance quickly is insufficient to outweigh the additional cognitive effort, especially as implicit adaptation should gradually improve performance over a relatively short period of training (Schween et al., 2020; Smith et al., 2006).

Evidence for explicit strategies in force-field adaptation has largely come from relatively simple tasks involving a single target, and only a handful of studies have examined the accompanying changes in preparation time. As the number of targets increases, discovering an underlying rule that supports an algorithmic strategy may become more difficult, and implicit adaptation may substantially reduce errors before such a strategy can be developed. In visuomotor rotation tasks, by contrast, the relatively simple relationship between target and aiming directions makes an algorithmic strategy a viable solution, especially when the set size is large. If participants instead learn separate force-field responses for individual targets, working-memory limitations may constrain strategy use. In visuomotor rotation tasks, the capacity to cache strategies consisting of individual stimulus–response associations appears limited to approximately three or four items, similar to spatial working memory (Velázquez-Vargas & Taylor, 2024), and performance deteriorates when the number of associations exceeds this capacity without an underlying structure to support an algorithmic strategy (Bejjanki & Taylor, 2026). Whether participants can similarly cache force-field strategies, and how many associations they can retain remains unknown. Caching a force-field strategy could impose greater demands on working memory because it may require retaining a time-varying force profile and the benefit of these strategies may diminish rapidly as implicit adaptation accumulates. Future work will be needed to estimate working memory capacity constraints for force-field strategies.

An unexpected finding was that the relationship between preparation time and force magnitude was steeper for incongruent targets. We included two targets, whose locations could switch within a trial pair, to limit planning before the target onset. We anticipated that switching targets would impose a fixed preparation time cost, increasing the intercept without changing the slope. Instead, we found that there was a steep increase in the slope when transferring the strategy to a different target. One possible explanation is that congruent and incongruent trials required different amounts of explicit compensation. When the target location is repeated, implicit adaptation should reduce the required strategic compensation (Fine & Thoroughman, 2006). Given the relatively narrow width of generalization of implicit adaptation, this contribution would be virtually zero when the target switched. However, this account is unlikely for two reasons. First, participants compensated only partially, so it is unclear what determined the suboptimal degree of explicit compensation they selected and whether they adjusted it to offset implicit adaptation. Second, such an adjustment would require strategic planning to account for the implicit adaptation induced by the preceding trial. Interactions between explicit and implicit adaptation can arise because these processes operate in series (Day et al., 2016; McDougle et al., 2017; Chen & Taylor, 2026; Taylor & Ivry, 2011; Miyamoto et al., 2020). Here, however, an explicit strategy would need to anticipate an implicit update that has not yet been observed and may be indistinguishable from noise. Although the motor system appears to account for its own uncertainty (Ranjan & Smith, 2018; Kim et al., 2026), it remains unclear whether explicit planning can estimate and incorporate these single-trial updates. Participants’ frequent surprise at their own aftereffects casts doubt on this possibility, suggesting that implicit recalibration is not readily accessible to deliberate planning or conscious awareness.

A more likely alternative explanation is that the preparation time pattern reflects the representational format of the algorithm used to counteract the force-field. In visuomotor rotation tasks, explicit re-aiming is often conceptualized as an angular transformation, analogous to mental rotation, in which participants rotate their intended movement direction away from the visual target (Taylor et al., 2014; McDougle & Taylor, 2019). Whereas the computational demand of this strategy scales with angular displacement, the present results suggest a computation whose demand scales with the magnitude of the lateral perturbation. Recent work shows that when a visuomotor mapping can be solved through either rotation or sign reversal, participants often favor sign reversal (Ding et al., 2026). Similarly, because the forces in our task were predominantly lateral, participants may have represented compensation as reversing and scaling a lateral displacement rather than rotating a movement plan. Under this representational account, experiencing a rightward perturbation during an upward reach would establish a representation in the first quadrant of movement space. Compensating at the same target could involve transforming this representation laterally into the second quadrant. Compensating at the opposite target might instead require an additional transformation into the fourth quadrant before reversing and scaling the lateral component into the third quadrant. If participants implemented these transformations by traversing an internal spatial representation, the additional path on incongruent trials could grow with perturbation magnitude, producing the steeper preparation time slope. This account is analogous to findings from mental imagery, in which the time required to scan between locations increases with their separation on an imagined map (Kosslyn et al., 1978), and potentially to magnitude-dependent processing times in mental arithmetic (Dehaene, 1992). However, this proposed sequence of transformations is highly speculative, and the present behavioral paradigm cannot uniquely identify the underlying representation. Ultimately, establishing the geometry of such cognitive representations for strategies may prove even more challenging than distinguishing the relatively straightforward debate concerning intrinsic and extrinsic reference frames (Brayanov & Smith, 2012; Poh & Taylor, 2019; Poh et al., 2026). Combining behavioral manipulations that distinguish candidate transformations with analyses of neural population trajectories may be required to reveal how these strategies are represented and implemented (Panichello & Buschman, 2021).

In summary, explicit force-field compensation incurred a preparation time cost that increased with perturbation magnitude and was further amplified when the response had to be transferred to a different target location. While these findings bear remarkable similarity to the algorithmic process supporting explicit re-aiming in visuomotor rotation tasks, its greater preparation time demands and sensitivity to target changes suggest that it may be more computationally demanding and operate on a different representational format. Together, these findings show that compensation for altered movement dynamics can recruit flexible, computationally costly explicit planning alongside implicit adaptation.

## Acknowledgements

We thank members of the Intelligent Performance and Adaptation Laboratory (IPA Lab) in the Department of Psychology at Princeton University for many fruitful discussions, and Samuel McDougle for discussions on representational geometries. This project was supported by grant R01NS131552 from the National Institute of Neurological Disorders and Stroke (NINDS) of the National Institutes of Health (NIH). HK, YW, and JAT were supported by grants R01NS131552 and R01NS134754 from the National Institute of Neurological Disorders and Stroke (NINDS) of the National Institutes of Health (NIH). The funders had no role in study design, data collection and analysis, decision to publish, or preparation of the findings.

## Notes

### Competing Interest Statement

The authors have declared no competing interest.

